# ADM: Adaptive Graph Diffusion for Meta-Dimension Reduction

**DOI:** 10.1101/2024.06.28.601128

**Authors:** Junning Feng, Yong Liang, Tianwei Yu

## Abstract

Dimension reduction is ubiquitous in high dimensional data analysis. Divergent data characteristics have driven the development of various techniques in this field. Although individual techniques can capture specific aspects of data, they often struggle to grasp all the intricate and complex patterns and structures. To address this limitation, we introduce *ADM (Adaptive graph Diffusion for Meta-dimension reduction)*, a novel meta-dimension reduction method grounded in graph diffusion theory. ADM integrates results from diverse dimension reduction techniques to leverage the unique strength of each individual technique. By employing dynamic Markov processes, ADM simulates information propagation for each dimension reduction result, thereby transforming traditional spatial measurements into dynamic diffusion distances. Importantly, ADM incorporates an adaptive mechanism to tailor the time scale of information diffusion according to sample-specific attributes. This improvement facilitates a more thorough exploration of the dataset’s overall structure and allows the heterogeneity among samples.

## 1 Introduction

Dimension reduction plays a crucial role in visualizing and comprehending complex datasets [1, 2]. The techniques in this field aim to condense high-dimensional data into a lower-dimensional representation while preserving the data’s inherent structure as faithfully as possible. This process is crucial for uncovering inherent similarities and differences between data points, thereby facilitating an intuitive understanding of the underlying patterns and trends [3, 4]. These techniques have become indispensable for exploratory analysis and pattern discovery across various domains, including computer vision [5], molecular biology [6], and genomics [7, 8], especially in single-cell transcriptomics [9–11].

Over the past few decades, researchers have explored a lot of dimension reduction techniques to reveal the structure of data characterized by noisiness and high dimensionality. Initial techniques, including Principal Component Analysis (PCA) [12], Multidimensional Scaling (MDS) [13], Sammon mapping [14], Isomap [15], and kernel PCA (kPCA) [16] have shown efficiency in delineating data with linear or non-linear characteristics by preserving dominant structures, while often at the expense of local details. Conversely, methods such as Locally Linear Embedding (LLE) [17], t-SNE [18], LargeVis [19], and Laplacian Eigenmaps [20] have prioritized the revelation of local structures but may sacrifice the global coherence of data. Subsequent developments, such as UMAP [2] and Hessian Locally Linear Embedding (HLLE) [21], have aimed at striking a balance between preserving both global and local structures. However, their effectiveness is sometimes compromised by sensitivity to noise and outliers. Furthermore, Diffusion maps [22] utilize a diffusion operator to explore the data’s intrinsic geometry, reducing the impact of noise, but facing challenges in preserving multiscale structures [23]. PHATE [24] advances the concept of the diffusion operator by integrating manifold learning and information geometry, providing a balanced representation of both global and local structures.

Since different techniques are developed based on distinct principles and paradigms, each is tailored to specific characteristics of the data. They often struggle to provide a comprehensive data representation, as they may overlook some other important characteristics of the data. In addition, each method involves hyperparameters that require fine-tuning for specific datasets, and different parameter configurations can lead to vastly different outcomes. Furthermore, due to the noisiness and high dimensionality that are prevalent in real-world datasets, their low-dimensional representations inevitably contain distortions from the underlying true structures, which can vary across different techniques.

Therefore, there is extensive interest in developing meta-analysis techniques capable of integrating the results from various individual methods, leveraging their strengths to achieve a more robust and comprehensive representation, and hopefully suppressing distortions in individual methods. Some methods, like fuzzy consensus analysis [25] and supervised learning methods [26–30], primarily focus on combining classification results from multiple datasets. However, meta-dimension reduction technologies require establishing a common space to align different results of the same data accurately. This necessitates further exploration and analysis of the intrinsic relationships between results from various methods. Recently, Ma et al. introduced *Meta-Spec* [31], which quantifies the relative performance of individual techniques in preserving the structure around each data point in the Euclidean space, and generates consensus dimension reduction results. This method has improved quality compared to individual techniques in capturing the underlying structure, providing more stable and superior dimensional reduction visualization results. However, it mainly offers a static measure that reflects the direct or physical distances between samples in Euclidean space, thereby facing challenges in capturing the dynamic geometric interactions and the complex mechanisms of information transmission among samples.

To address these challenges, we propose *ADM*, a novel meta-dimension reduction and visualization technique based on information diffusion. For each individual dimension reduction result, *ADM* employs a dynamic Markov process to simulate the information propagation and sharing between data points. It introduces an adaptive mechanism that dynamically selects the diffusion time scale based on the local context of individual samples, enhances the model’s ability to explore the multi-scale structures, and effectively mitigates the impact of noise. This process transforms the traditional Euclidean space dimension reduction results into an information space, thereby revealing the intrinsic manifold structure of the data. By leveraging the strengths of multiple dimension reduction techniques, *ADM* significantly improves the robustness of the dimension reduction result, making it a powerful tool for analyzing high-dimensional and noisy datasets.

## 2 Methods

### 2.1 Overview of the method

For each output from an individual candidate dimension reduction technique, the *ADM* approach initiates by transforming the Euclidean distance of features into a diffusion operator *𝒫*. This operator quantifies the likelihood of information propagation between samples through random walks. Subsequently, a sample-specific diffusion process is implemented for each sample to simulate multi-step diffusion. We employ the Breadth-First Search (BFS) algorithm to adaptively select the appropriate diffusion time scale for each sample. This adaptive strategy considers the inherent heterogeneity within datasets, effectively filters out noise and prevents over-smoothing. The resulting sample-specific propagation probabilities are then utilized to calculate the diffusion distances. These diffusion distances leverage the dynamic Markov process to link the Euclidean distance with geometric densities, highlighting the intrinsic similarities and differences among samples in the information manifold, serving as a robust metric for quantifying the relative positions within the information space [32].

Next, we combine the diffusion distance matrices from all candidate methods using harmonic averaging and gamma distribution-based normalization to construct a comprehensive meta-diffusion-distance matrix. This distance leverages the advantages of individual candidate techniques, providing a robust representation of the dataset and enabling in-depth exploration of complex relationships among samples. For visualization and further interpretation, the meta-distance matrix can be reduced to a low-dimension embedding via UMAP, producing plots that reveal the dataset’s underlying structure. The overall framework of *ADM* is shown as Figure 1.

**Fig. 1.**
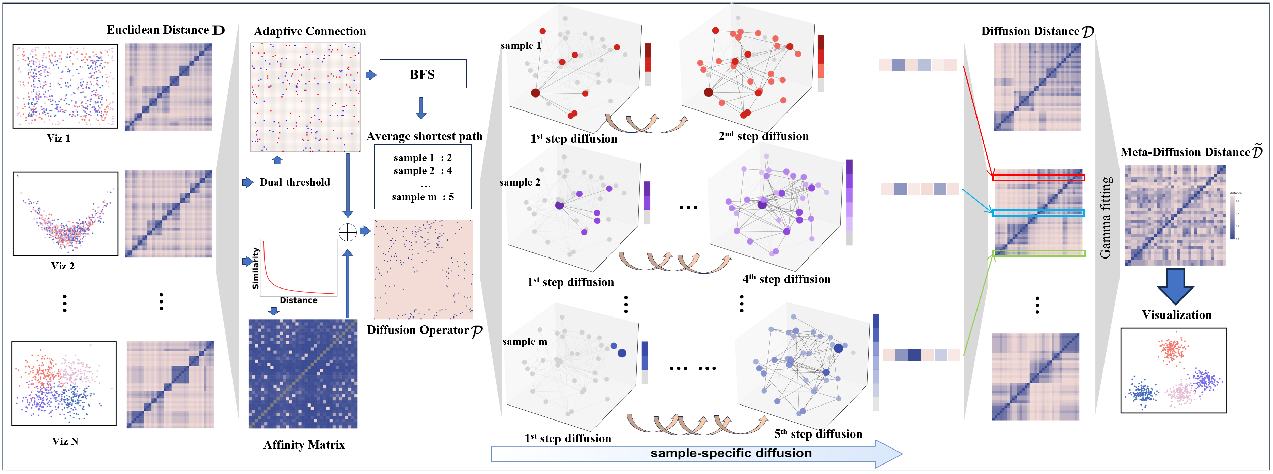
The framework of *ADM*. It takes inputs from various dimension reduction techniques and generates a meta-dimension reduction and visualization output that integrates the strengths of these individual results.

### 2.2 Data preprocessing

In our method, the input data comprises a set of feature matrices generated by various dimension reduction techniques (*N* × *𝒳, 𝒳* ∈ ℝ ^*m*×*n*^), known as candidates, where *N* is the counts of candidates, *m* denotes the total number of samples, and *n* indicates the feature dimension. Initially, we preprocess each matrix *𝒳* to move outliers potentially skewing the dataset’s true structure, thereby preserving the data integrity and consistency. Specifically, outliers are first discovered by the method by Knorr and Ng [33].

For each feature ***x***_**·*k***_ (i.e., the *k*^*th*^ column) in *𝒳*, we calculate the interquartile range *IQR* = *Q*_3_ − *Q*_1_, where *Q*_3_ is the 75^*th*^ percentile and *Q*_1_ is the 25^*th*^ percentile of ***x***_**·*k***_. We set the neighborhood radius threshold to *d* = *IQR/*3 for the method by Knorr and Ng [33]. We remove all data points detected as outliers from ***x***_**·*k***_ to generate a new vector 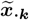 Our objective is to adjust the original ***x***_**·*k***_ to resemble the distribution of 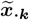 To achieve this goal, we employ quantile normalization of ***x***_**·*k***_ using 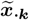 as the reference distribution. The effect of this quantile normalization is the outliers in the feature dimension shrink towards the data cloud, while non-outlier data points are changed very little.

### 2.3 Affinity matrix

After preprocessing, we first calculate the Euclidean distance matrix D, in which *d*_*ij*_ is the Euclidean distance between sample *i* and *j*. To eliminate self-comparisons, we set *d*_*ii*_ as infinity. Subsequently, we convert the distance matrix **D** into a affinity measure **S**:

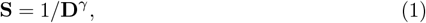

where *γ* is the parameter for the power transformation, and **S** signifies the sample affinities in the Euclidean space. This non-linear scaling mitigates the overwhelming impact of larger distances, thus emphasizing local neighborhood structures.

### 2.4 Adaptive connectivity

In information diffusion theory, connectivity between nodes is crucial for mapping out the routes and patterns of information propagation. To accommodate the complexity of the relation between samples and capture multi-scale structure, we introduce a two-step procedure to construct a sample connectivity matrix **A** based on the distance matrix. This method considers both the overall trends and patterns across the entire dataset, as well as the local context surrounding individual samples. Globally, we set a global percentile threshold *α* to identify crucial sample interactions across the dataset. For example, setting *α* = 5%, designates the shortest 5% of distances in **D** as globally significant, with *thres* = *percentile*(**D**, *α*). If *d*_*ij*_ ≤ *thres*, the connection between samples *i* and *j* is considered substantial on a global scale, indicated by **A**_*ij*_ = 1; otherwise, **A**_*ij*_ = 0.

Locally, the percentile threshold *β* is used to adjust the connectivity of each sample according to its immediate neighbors, with *thres*_*i*_ = *percentile*(***d***_*i*·_, *β*) determining local connections, where ***d***_*i*·_ represents distances from sample *i* to all others. If *d*_*ik*_ ≤ *thres*_*i*_, (0 *< k* ≤ *n, k*≠ *i*), we set **A**_*ik*_ = 1. Using the two-step procedure to construct the connectivity matrix helps to strike a balance between global exploration and local structure sensitivity.

To establish sample-specific affinity relations, we further generate a sparse adaptive affinity matrix (*U*) by applying an element-wise multiplication between the affinity matrix **S** and the connectivity matrix **A**:

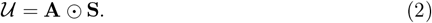

This operation effectively strengthens the connections between samples of high similarities while diminishing the influence of those between dissimilar samples. In addition, this affinity measure only accounts for the local distance between samples. Ultimately, we transform these weighted connectivities to a weighted undirected graph ***𝒢*** = {*V, E*}, where *V* represents individual samples, and *E* denotes the connections defined by matrix **A**, with edge weights given by *𝒰*.

### 2.5 Sample-specific diffusion

We use the Markov random walk process to explore the nonlinear manifold structure of the dataset. The transition probabilities in this random walk are calculated through row-normalization of matrix *𝒰*, which renders the adaptive affinity into a Markov-based diffusion operator *𝒫*:

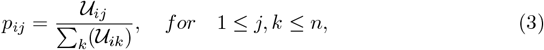

where *p*_*ij*_ ∈ *𝒫* denotes the probability of information propagating from sample point *i* to point *j*. We refer to *𝒫* as the diffusion operator, which quantifies the likelihood and intensity of information propagation across the weighted graph.

The time scale *τ*, depicting the depth of information propagation in a random walk, plays a crucial role in balancing the trade-off between capturing local details and global information. A smaller value of *τ* emphasizes local structures, often associated with noisy information, while a larger *τ* captures global structures but may smooth out important details. Hammond et al. [34] pointed out that nodes within a densely connected graph may exchange information more efficiently than their sparsely connected counterparts. Therefore, our approach adaptively selects the optimal *τ* for each sample by evaluating its spatial position and connectivity to ensure a balanced representation of local and global characteristics.

To achieve this goal, we initially compute the shortest path lengths across all sample pairs within graph ***𝒢***. We record the results in a matrix **L** ∈ ℝ ^*m*×*m*^, where *m* is the total number of samples, and each element *l*_*ij*_ ∈ **L** denotes the shortest path length from node *i* to node *j*.

We then calculate the average path length for each sample as a metric of its connectivity within the overall graph:

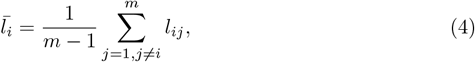

where 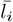 denotes the average shortest path length for sample *i* relative to others.

The diffusion steps of sample *i* is then set to a value that is proportional to its average steps,

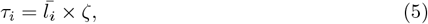

where ζ is a pre-determined scaling constant. By employing this approach, we ensure the diffusion process is finely tuned to reflect the unique connectivity and structural properties of each sample. Samples on the edge of the point cloud are allowed more diffusion steps, while samples in the center use fewer diffusion steps. This adaptive process enhances the model’s sensitivity to local and global connectivity patterns, facilitating a more precise depiction of the complex distribution patterns within datasets.

Next, we calculate the diffusion distance of each sample relative to other samples by incorporating the previously determined diffusion steps {*τ*_*i*_}_*i*=1,…,*m*_ specifically selected for each sample with the Markov-based diffusion operator *𝒫*. Specifically, the diffusion distance from sample *i* to *j* after *τ* steps of a random walk is calculated as follows:

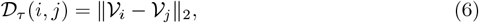

where 𝒱_*i*_ = − log(***p***(*τ*_*i*_|*i*)), *i* = {1, 2, …, *m*}, and ***p***(*τ*_*i*_|*i*) represent the likelihood of information propagation from sample *i* to the rest of the graph after *τ*_*i*_ steps of diffusion. Here, ***p***(*τ*_*i*_|*i*) ∈ *𝒫* (*τ*_*i*_) and 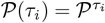 represent how samples are interconnected through a *τ*_*i*_-step diffusion process. Thus, the computation of diffusion distance serves as a bridge that smoothly transitions from local statistical properties to global manifold structures. The computation of the graph diffusion process is intensive. In actual computation, to save computing time, we first calculate the diffusion for all samples using a set of quantiles of all {*τ*_*i*_}_*i*=1,…,*m*_, and then for sample *i*, we use the diffusion results obtained from the quantile that *τ*_*i*_ is closest to.

Subsequently, we symmetrize the diffusion distance matrix to guarantee that the distance from any sample to another is reciprocally consistent:

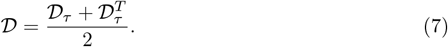

*𝒟* represents the diffusion distance matrix associated with a specific dimension reduction result. This sample-specific computation of diffusion distances yields a multi-scale family of information geometry representations across the samples, offering a detailed perspective on overall patterns and heterogeneities.

### 2.6 Normalization and Integration

After applying the above procedure to each individual dimension reduction result, we obtain a set of diffusion distance matrices { *𝒟*_*k*_}_*k*=1,…,*N*_. An important aspect of our approach involves integrating diffusion distance matrices derived from various candidates. For this purpose, we employ a two-step procedure involving distribution normalization and matrix integration.

#### Distribution normalization

The diffusion distance matrices contain all positive values with heavy right-skewness. To suite the data, we employ Gamma distribution to fit the obtained diffusion matrices. The probability density function of the Gamma distribution is represented as:

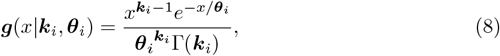

where 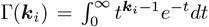 denotes the Gamma function. We first determine the shape parameter ***k***_*i*_ and scale parameter ***θ***_*i*_ of the Gamma distribution for each diffusion distance matrix *𝒟* _*i*_ using standard maximum likelihood estimation (MLE) procedure:

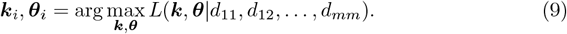

With the *MLE* method, we determine the shape and scale parameters for each diffusion distance matrix to capture inherent variability across different dimension reduction techniques. Subsequently, we compute the average parameters 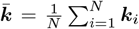 and 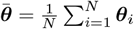 across all matrices, reflecting the aggregate distribution features.

Then we construct a reference distribution ***O*** using gamma distribution with the averaged parameters 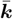 and 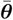, and employ quantile normalization to align each of the diffusion distance matrices within { *𝒟*_*k*_}_*k*=1,…,*N*_ with the reference distribution ***O*** to obtain the corresponding normalized distance matrices 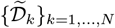, using the computation method described in Section 2.2. This normalization ensures that the distributions of the diffusion distance are comparable with each other, facilitating meaningful comparisons and mergers between diffusion distance matrices. The final meta-diffusion distance matrix is obtained by exponentiating and summing the candidate normalized matrices as follows:

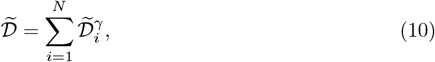

where *γ* serves as a power parameter that tunes the relative influence o f l arge and small distance values. The meta diffusion distance matrix is then utilized to generate a meta-visualization with established techniques such as UMAP or PCA.

## 3 Results

### 3.1 Evaluation criteria

To thoroughly evaluate the performance of dimension reduction models, this study uses various evaluation metrics to assess these techniques from multiple perspectives.

#### Category consistency index (*CCI*)

This indicator is introduced to assess the models’ capability to preserve original data category similarities despite the presence of noise. The *CCI* is a metric used to evaluate the consistency between the distances among samples and their respective categories after dimension reduction. Inputs for *CCI* include a distance matrix 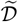, obtained from dimension reduction, and the true category labels of the samples. Specifically, *CCI* is computed by identifying the nearest top-*r* samples for each dataset sample and calculating the average proportion of those that share the same category, yielding a value between 0 and 1. A *CCI* value closer to 1 indicates that the distances obtained through a dimension reduction technique more effectively reflect the true categorical relationships between samples in its local structure, making the distances more meaningful. Hence, *CCI* serves as an informative metric for evaluating the efficacy of distance representations. In the current study, we use three variants of this indicator, *CCI*_*raw*_, *CCI*_*umap*_, and *CCI*_*pca*_, which measures the preservation of category similarity within the raw distance matrix, as well as the feature matrices after dimension reduction by UMAP and PCA, respectively.

#### Structural consistency index (*SCI*)

The *SCI* metric evaluates the structural fidelity of data after dimension reduction relative to the original, noise-free structure:

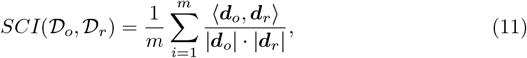

where *𝒟* _*o*_ and *𝒟* _*r*_ denote the distance matrices of the original data and after dimension-reduction, respectively, ***d***_*o*_ and ***d***_*r*_ are row vectors in these matrices, and *m* is the total number of data points. This indicator falls into [0, 1]. A higher *SCI* value signifies higher concordance between the origin data structure and the one after a dimension reduction.

#### Adjusted Rand Index (*ARI*)

The *ARI* metric evaluates the similarity of clustering results to true labels. It produces values within the range of [−1, 1], where a score close to 1 indicates high agreement with true labels, while a score around 0 indicates no better than random label assignment, and a score close to -1 implies dissimilarity to true label distribution.

#### Normalized Mutual Information (*NMI*)

This indicator quantifies the mutual information between clustering outcomes and true labels, normalized to fall within [0, 1]. Higher *NMI* values indicate greater alignment between clustering results and true labels, with values approaching 0 indicating minimal consistency.

### 3.2 Simulation studies

In this section, we generate a large amount of representative simulation data to assess the universal applicability and robustness of dimension reduction techniques. We conduct simulations from two different perspectives. In Section 3.2.1, we compare the ability of different meta-dimension reduction technologies to mitigate interference from low-quality outputs by simulating the output of multiple reduction techniques at different quality levels. In Section 3.2.2, we generate several original data with typical structures and compare the ability of different dimension reduction techniques to restore the underlying structure of the data by adjusting signal-to-noise ratios.

#### 3.2.1 Simulation on synthetic datasets

Dimension-reduction outputs from various candidate methods often range from high to low quality for a given dataset. Traditional integration methods may struggle with these imbalanced inputs, potentially skewing integrated results towards lower-quality inputs. This simulation assesses the meta-visualization technologies’ ability to discern and prioritize high-quality outputs over low-quality outputs. Such distinctions are crucial for evaluating the meta-visualization technologies’ effectiveness in extracting meaningful patterns from candidate results.

##### High-quality outputs

Firstly, we generate three sets of independent data points ***x***_01_, ***x***_02_, ***x***_03_ ∈ ℝ ^*n*×2^ representing three types of samples, respectively. These sets initially share identical means and covariance matrices. Then, in each candidate dimension reduction result, we introduce differences between types of samples by applying random rotational and stretching transformations to each set:

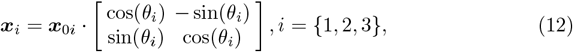

where the angle *θ*_*i*_ is randomly chosen from [0, 2*π*]. Subsequent, we apply scaling and translational transformations to ***x***_1_ and ***x***_2_:

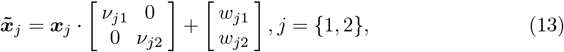

here, *ν*_*j*·_ ∈ ℝ are scaling factors, *w*_*j*·_ ∈ ℝ are translational transformation factors. 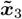 stays the same as ***x***_3_. The transformed sets 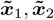, and 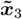 are concated as a simulated high-quality candidate output ***y***_1_ ∈ ℝ^3*n*×2^. This procedure is replicated *p* times to produce the candidate high-quality outputs ***s*** = {***y***_*i*_}_*i*=1,…,*p*_ ∈ ℝ ^3*n*×2^.

##### Low-quality outputs

To simulate poorly-performing outputs, we generate *q* sets of random noise data ***z*** ∈ ℝ^3*n*×2^. Each set is generated with a specified mean *µ*_1_, *µ*_2_ ∈ ℝ and variance-covariance matrix **Σ**_*k*_:

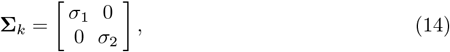

where *σ*_1_, *σ*_2_ ∈ ℝ represent the variance on each dimension. The generation process is iterated *q* times to create diverse low-quality outputs ***z*** = {***z***_*i*_}_*i*=1,…,*q*_ ∈ ℝ^3*n*×2^, integrated with the high-quality data **s** to construct comprehensive inputs of meta-dimension reduction methods. In our simulation, we set *n* = 200 as the sample number of each type, the scaling factor *ν*_1_ = 1, *ν*_2_ = 4, the translational factors *w*_1_ = 0.5, *w*_2_ = 2, and signal-containing data counts *p* = 3. Additionally, we randomly select ten different numbers from 1-100 to simulate various counts of low-quality outputs and generate such data with parameters *µ*_1_ = 1, *µ*_2_ = 12, *σ*_1_ = 0.1 and *σ*_2_ = 15.

Figures 2(A-C) present a comparative analysis of the Category consistency index (CCI) scores achieved by different meta-dimension reduction methods when faced with varying quantities of low-quality sets within the candidate outputs. The experiments were conducted with different values of *r* = {1, 2, 5, 10, 20}. Specifically, Figure 2A illustrates the *CCI* scores calculated directly using the distance matrix output from the *ADM* method and the meta-distance matrix from the *Meta-Spec* method. Figure 2B and Figure 2C demonstrate the results after applying dimension reduction to the distance matrices using UMAP and PCA, respectively. The results indicate that while both *ADM* and *Meta-Spec* experience a decrease in *CCI* scores with an increase in low-quality group sets, *ADM* consistently achieves higher *CCI* scores than *Meta-Spec* across various levels of noise.

**Fig. 2.**
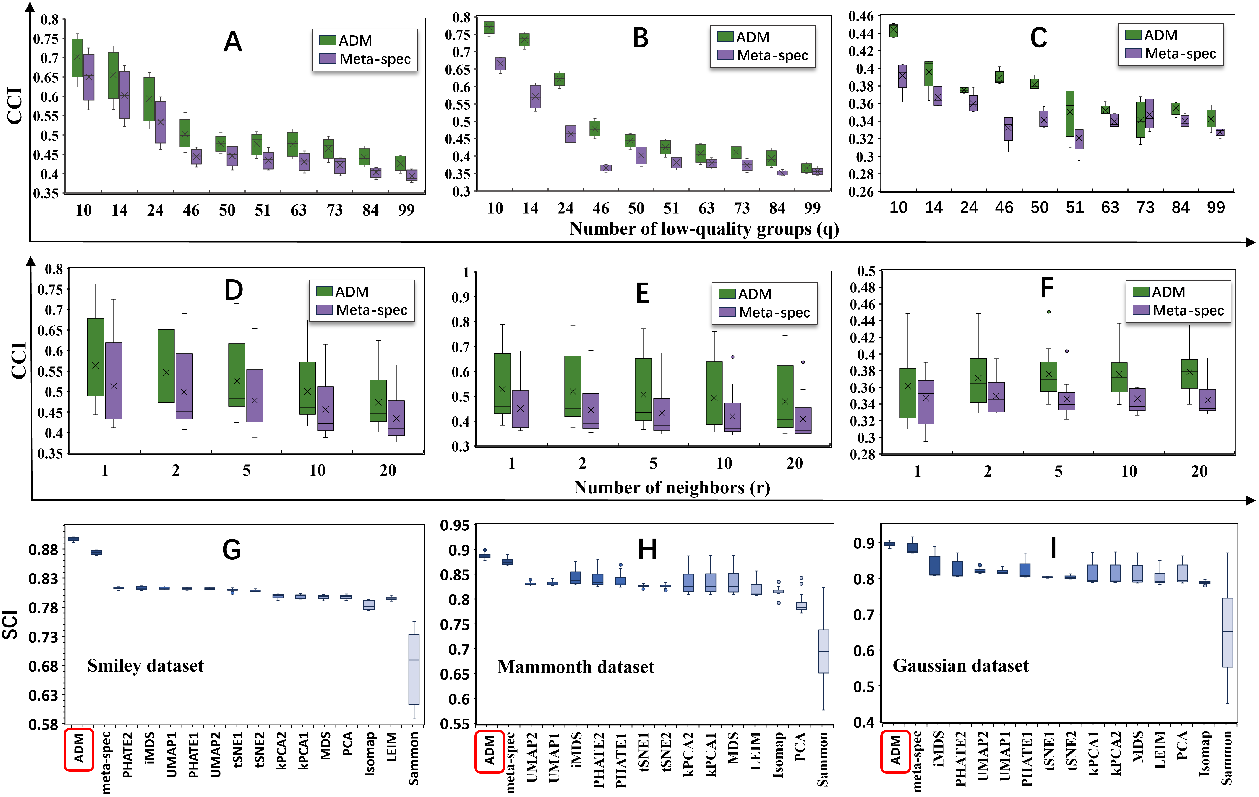
Comparative Performance of *ADM* and *Meta-Spec* Methods on Simulated Data. **A-F:** Category Consistency Index (*CCI*) values of *ADM* and *Meta-Spec* on simulation data of different dimension-reduction outputs. **A-C:** The *CCI*_*raw*_, *CCI*_*umap*_, and *CCI*_*pca*_ under different noise conditions, given fixed ranking parameters *r*. **D-F:** Giving different groups of noise, the *CCI*_*raw*_, *CCI*_*umap*_, and *CCI*_*pca*_ under varying ranking parameters. **G-I:** The 15 comparative results of the Structural Consistency Index (SCI) across varying signal strengths for the Smiley Dataset, Mammoth Dataset, and Gaussian Dataset, respectively. HLLE is omitted due to very low consistency.

Figures 2(D-F) display the *CCI* values obtained by the two meta-dimension reduction methods at different r values, ranging from 1 to 20. Simultaneously, we randomly selected ten numbers from the range of 1 to 100 as the number of low-quality sets. It is observed that the *CCI* values for both *ADM* and *Meta-Spec* decrease as r increases, which is expected as the value of *r* determines how many neighbors are considered. However, it is evident that *ADM* outperforms *Meta-Spec* in terms of *CCI*_*raw*_, *CCI*_*umap*_, and *CCI*_*pca*_ across all tested r settings.

These experimental findings directly reflect *ADM* ‘s superior capability in mitigating the impact of noise from low-quality candidate methods on the signal from high-quality methods. More fundamentally, since *CCI* measures the consistency between sample distances and category labels, the results essentially indicate that the integration of diffusion distance matrices in *ADM* better captures the distribution of the original data compared to the integration of Euclidean distance matrices in *Meta-Spec*.

#### 3.2.2 Simulation to test structure restoration

Dimension reduction serves a crucial goal of retaining essential and representative data features while mitigating complexity. In some specific fields, such as single-cell data analysis, high-noise environments pose significant challenges to dimension reduction techniques in extracting signals and preserving the original structure. Therefore, we generate several high-dimensional noisy simulation datasets to evaluate the effectiveness of dimension reduction techniques in reconstructing original data structures and mitigating noise interference.

Specifically, we generate this simulation data by employing a signal-plus-noise model: ***Y*** = ***Y*** ^*^ + ***Z***, where ***Y*** ∈ ℝ ^*n*×*p*^ is the high-dimensional observations, ***Y*** ^*^ represent the noise-free signals that inherently reside within an *h*-dimensional linear subspace, derived from varied low-dimensional structures. However, they are subject to an arbitrary rotation within the ℝ ^*p*^ space, resulting in them being represented as *p*-dimensional vectors with dense (nonzero) coordinates. ***Z*** ∈ ℝ^*n*×*p*^ represents random noise sampled from a standard multivariate normal distribution. This approach enables us to controllably model the intricate structures present in noisy observational data. We construct three datasets with different low-dimensional original structures. **Smiley Dataset (*h*** = 2): Derived from a two-dimensional smiley face configuration, this dataset replicates complex nonlinear structures commonly observed in real-world data. **Mammoth Dataset (*h*** = 3): Comprising points from a three-dimensional mammoth structure, this dataset simulates three-dimensional forms within a high-dimensional space. **Gaussian Mixture Dataset (*h*** = 6): Consisting of six categories of observations, this dataset allows for the exploration of mixed signals in high-dimensional domains.

In the experiment, we introduce varying noise levels by adjusting the signal-to-noise (SNR) ratio *φ* to simulate realistic high-dimensional, high-noise data. Specifically, we randomly select 20 values of *φ* from [0.0001, 1] to generate a series of noise ***Z***_*i*_, *i* = 1, …, 20 for each dataset. For the **Smiley** dataset 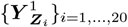, we set the total number of observation points *n* as 550 and the feature dimension *p* as 300. Similarly, for the **Mammoth** 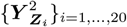 and **Gaussian Mixture** dataset 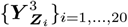, we set *p* to be 300, and 500, respectively and *n* at 1000, and 900, respectively.

Then, we apply 11 individual dimension reduction techniques, including PCA, UMAP, PHATE, etc, and a meta-dimension reduction technique *Meta-Spec* as comparative methods. We test some of these individual techniques with 2 different parameter settings, and finally obtained 15 individual candidates. *ADM* and *Meta-spec* use these 15 candidates as inputs. We evaluate the performance of *ADM*, against these 15 comparative methods using the Structure Consistency Index (*SCI*) metric.

Figure 2 (G-I) displays the performance comparison of 16 different methods applied to three distinct datasets 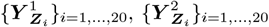 and 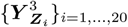 respectively. Each boxplot shows the distribution of the achieved *SCI* values under 20 different levels of noise. The results show that meta-visualization techniques, including *ADM* and *Meta-Spec*, outperform individual dimension reduction methods in maintaining the core structure of the data. Across the three data sets, the *SCI* values of *ADM* are higher than those of *Meta-Spec*. Notably, *ADM* is characterized by narrower boxplots, which is more pronounced in the Mammoth and Gaussian Mixture datasets, indicating its superior consistency across datasets with different levels of noise. This highlights *ADM* ‘s strong capacity for noise reduction, signal detection, and accurate reconstruction of the data’s original structure through its dynamic information diffusion mechanism.

### 3.3 Results on Real Data

In this section, we apply the proposed *ADM* method together with *Meta-spec* and 15 individual dimension reduction methods to six publicly available single-cell datasets to evaluate its performance in dimension reduction and visualization. These datasets encompass single-cell RNA sequencing data from the mouse midbrain and striatum, human lymphocytes, mouse tendon cells, a mixed cell population from the human immune and respiratory systems, and the single-cell RNA-seq and mirRNA-seq data from human cancer cell lines.

For each dataset, we begin by applying 11 individual dimension reduction techniques under different parameter settings, resulting in 15 candidates and saving their outputs. Subsequently, we utilize meta-dimension reduction methods, including *ADM* and the comparison method *Meta-Spec*, to integrate these results. The overall accuracy in terms of preserving between-cell type differences based on true cell labels is summarized by ARI and NMI, which are calculated after UMAP dimension-reduction of the respective distance matrices (Figure 3). To avoid artifacts caused by clustering techniques, we also analyzed the preservation of true cell class information in the distance matrices before and after further dimension reduction (Figure 4). We discuss the details of these results in each of the real datasets in the following sub-sections.

**Fig. 3.**
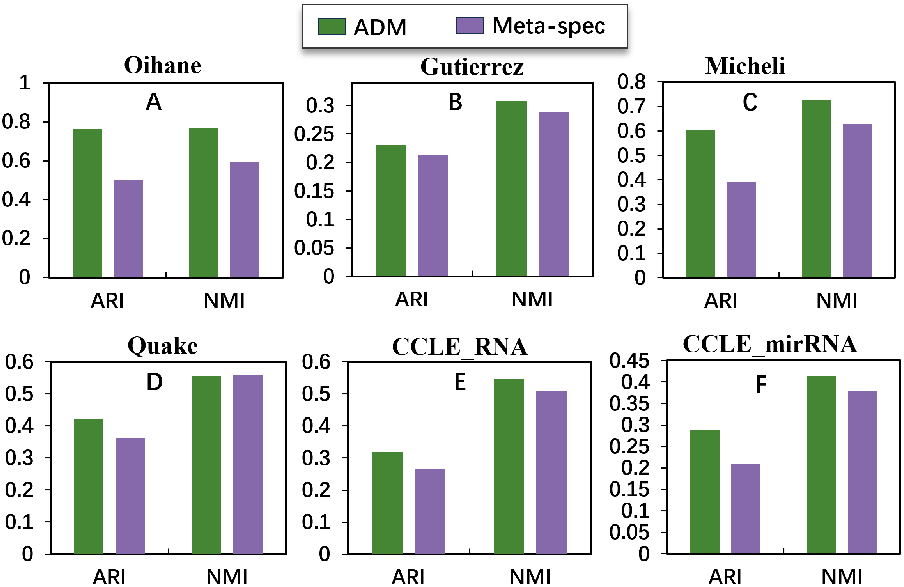
The *ARI* and *NMI* results on real data.

**Fig. 4.**
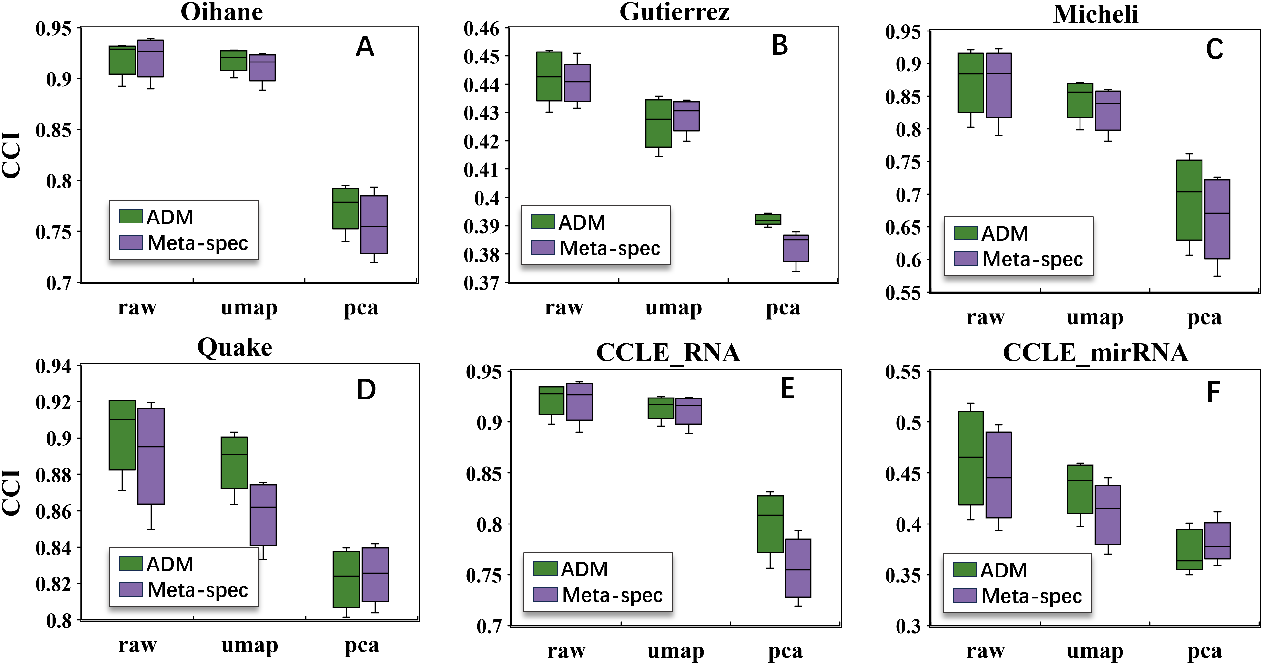
The *CCI* results on real data.

#### 3.3.1 Oihane dataset: mouse midbrain and striatum scRNAseq data

The first dataset is the single-cell RNA sequencing data of the mouse midbrain and striatum, named Oihane [35]. It comprises expression profiles from 1,331 single cells and 213 genes. This dataset covers various cell types, including oligodendrocytes, microglia, neurons, astrocytes, endothelial cells, ependymal cells, and hybrid cells.

Figure 5 shows the visualization outcomes of various individual dimension reduction techniques and two meta-reduction techniques. Among the individual techniques, UMAP and t-SNE excel in separating different categories, with t-SNE exhibiting poor intra-class compactness. PHATE, Isomap, and LEIM capture the global manifold structure of the data but are less effective in differentiating between cell types. In contrast, meta-dimension reduction techniques can leverage the strengths of individual reduction techniques to maintain proximity relationships among data points while revealing the overall manifold structure of the dataset.

**Fig. 5.**
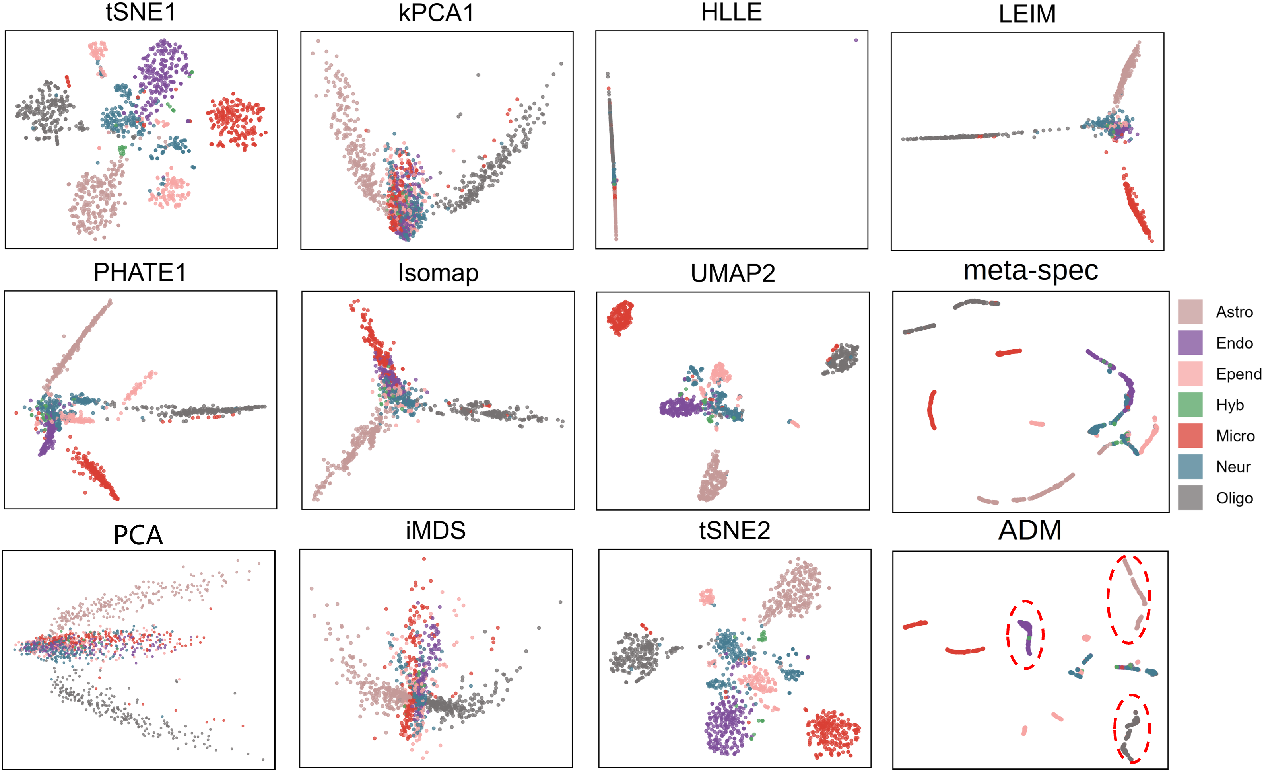
Visulization of the Oihane dataset. The abbreviations in the figure correspond to their full names: Astro: Astrocytes, Endo: Endothelial, Epend: Ependymal, Hyb: Hybrid, Micro: Microglia, Neur: Neurons, Oligo: Oligodendrocytes. Due to space limitations, we present the outcomes of 10 randomly selected individual techniques. The complete comparative results can be found in Figure S1 in the supplementary materials.

Compared with *Meta-Spec, ADM* further expands the distance between cell types while maintaining intra-class tightness. In the visualization, the distances produced by *Meta-Spec* tend to identify oligodendrocytes as two distinct categories, one of them overlapping with microglia, whereas the diffusion distance generated by *ADM* closely arranges oligodendrocyte cells and creates clear separation from other categories. For Endothelial and neurons, *ADM* exhibits a clear boundary, whereas in the *Meta-Spec* visualization, these two cell types overlap with blurred boundaries. Additionally, in *Meta-Spec*, some astrocytes intertwine with neurons in the visualization, while the *ADM* results form two distinct clusters with clear boundaries between them.

Additionally, Figure 3A demonstrates that *ADM* is superior in clustering accuracy, offering significantly higher clustering precision of Adjusted Rand Index (*ARI* of 0.7650) and Normalized Mutual Information (*NMI* of 0.7686), compared to the *ARI* (0.4999) and *NMI* (0.5963) of *Meta-Spec*. Furthermore, as Figure 4A illustrates, the *ADM* consistently outperforms *Meta-Spec* in preserving category similarities, suggesting that the adaptive diffusion distance produced by the proposed *ADM* can serve as a robust representation of cells for subsequent analysis.

#### 3.3.2 Gutierrez dataset: human lymphocyte population

The dataset is single-cell RNA sequencing (scRNA-seq) data from human lymphocyte populations [36]. It captures the gene expression of 662 genes across 2,036 single cells, including iNKT, MAIT, *λδ*T cells, NK cells, and CD4+ and CD8+ T cells, which demonstrated a high degree of functional and phenotypic similarity.

As shown in Figure 6, current individual dimension-reduction techniques such as UMAP, t-SNE, iMDS, and PHATE face significant challenges in distinguishing these subtypes. They struggle to segregate any one subtype distinctly or allocate these cells to appropriate categories.

**Fig. 6.**
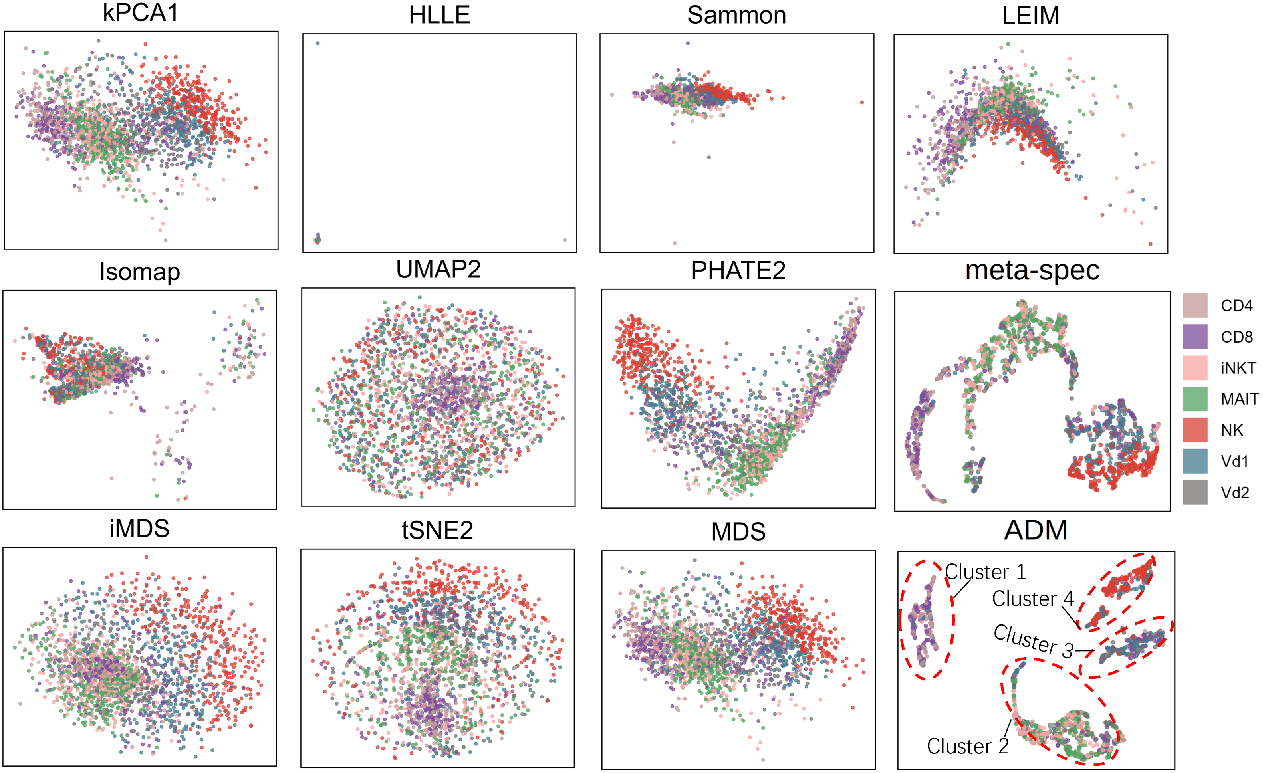
Visulization of Gutierrez dataset. The complete comparative results can be found in Figure S2 in the supplementary materials.

*ADM* can reveal subtle differences between cell subtypes. In the visualization analysis, we identified four clear clusters representing a continuum of lineages from adaptive to innate characteristics. Cluster 1 primarily consists of adaptive T cells (CD4+ and CD8+) expressing molecules related to antigen presentation and specificity recognition. Cluster 2 encompasses iNKT, MAIT, and V*δ*2 T cells, subtypes that express genes associated with rapid immune responses, reflecting their innate-like T cell features. Cluster 3 includes *λδ*1 and *λδ*2 T cells, which are characterized by gene expressions related to innate immune surveillance and cytotoxic functions. Cluster 4 is composed of NK cells, whose expression profiles are directly linked to innate immune responses.

Compared to the *Meta-Spec* method, which could partially distinguish some cell subtypes, *ADM* is notably clearer in terms of category boundary definition and inter-class distance. The ARI and NMI values of *ADM* are slightly higher than those of *Meta-Spec* (Figure 3). By leveraging the mechanism of information diffusion, *ADM* not only enhanced the accuracy of cell type identification but also provided new perspectives on the functional balance of immune cells in immunological defense, offering important insights for a deeper understanding of the complexity of the immune system and the development of targeted immune therapeutic strategies.

#### 3.3.3 Micheli dataset: tendon cell population

We analyzed RNA data from 1,191 single-cell samples within the mouse Achilles tendon, covering 3,809 genes and 13 distinct cell types. This dataset includes three subpopulations of tendon fibroblasts—Tendon Fibroblasts 1, Fibroblasts 2, and Connective Fibroblasts, as well as pericytes, immune cells, erythrocytes, and sheath fibroblasts. Additionally, there are three groups of endothelial cells and three different nerve cell populations, we call this dataset as Micheli [37].

In the *Meta-Spec* visualization (Figure 7), while clusters of cell types such as endothelial cells, immune cells, and nerve cells could be recognized, the separation within these categories was not clearly defined. Notably, some overlap between 16 endothelial cell subtypes and tendon fibroblasts was observed, suggesting potential limitations of *Meta-Spec* in discriminating closely related cell populations. In contrast, the *ADM* visualization presented much clearer boundary between cell types, particularly in distinguishing between subtypes of nerve cells and immune cells. Furthermore, *ADM* successfully achieved a distinct separation between erythrocytes and tendon fibroblasts.

**Fig. 7.**
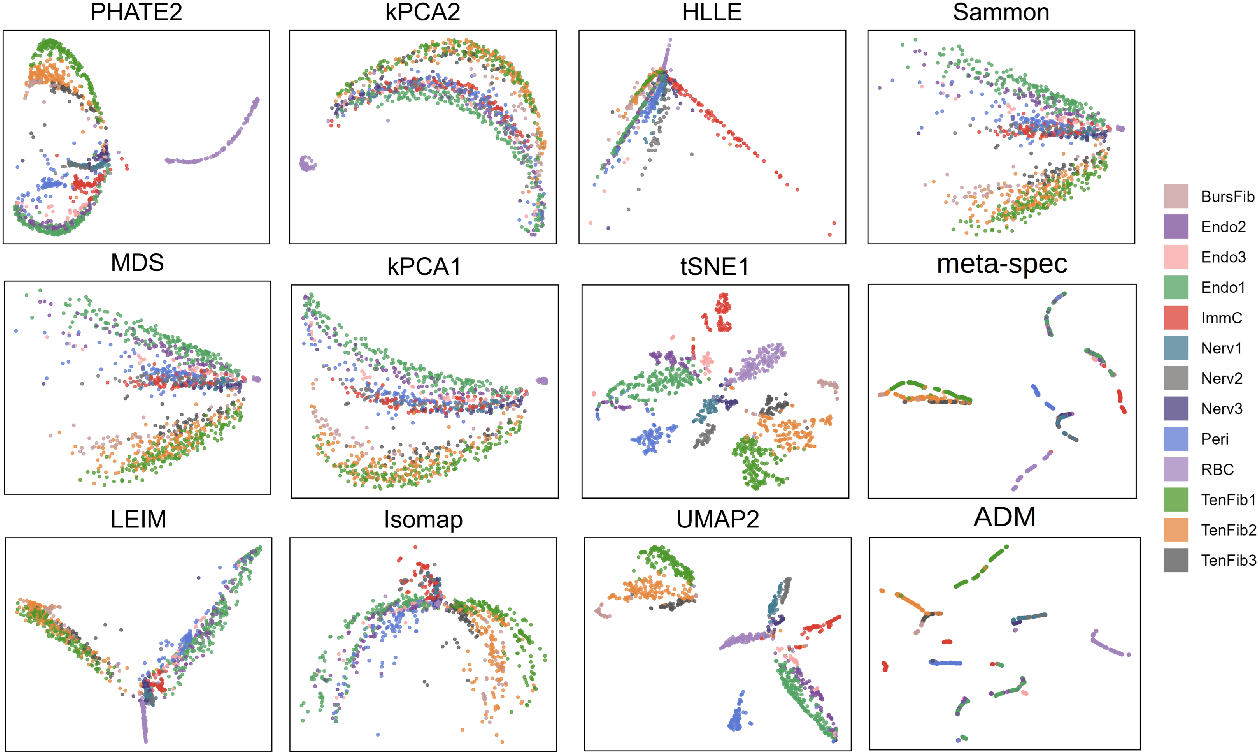
Visulization of the Micheli dataset. The abbreviations in the figure correspond to their full names: BursFib: Bursal fibroblasts, Endo: Endothelial cells, ImmC: Immune Cells, Nerv: Nerve cells, Peri: Pericytes, RBC: Red blood cells, TenFib: Tendon fibroblasts. The complete comparative results can be found in Figure S3 in the supplementary materials.

Quantitative metrics further supported these visual observations. *ADM* ‘s Adjusted Rand Index (*ARI*) of 0.6037 was significantly higher than *Meta-Spec*’s 0.3895. Likewise, *ADM* ‘s Normalized Mutual Information (*NMI*) of 0.7256, compared to *Meta-Spec*’s 0.6292, better reflected its efficacy in preserving the information within the dataset.

In summary, while *Meta-Spec* provided valuable data insights, *ADM* demonstrated superior performance in clustering purity and fidelity of data representation.

#### 3.3.4 Quake dataset: mixed cell types of human immune and respiratory systems

The dataset [38] consists of sequencing results for 3,349 genes across 1,676 single cells. It includes a diverse range of cell types, including stromal cells, lung epithelial cells, lung endothelial cells, B cells, leukocytes, monocytes, T cells, classical monocytes, ciliated columnar cells of the tracheobronchial tree, myeloid cells, and natural killer cells. These cell types represent various components of the human immune and respiratory systems.

Figure 8 shows the visualization of *ADM* and representative comparison methods. TSNE, LEIM, and UMAP demonstrated their ability to distinguish T cells and lung endothelial cells from other cell types with noticeable clarity. However, the intra-class aggregation of these individual methods is not as good as meta-dimension reduction methods *Meta-Spec* and *ADM*.

**Fig. 8.**
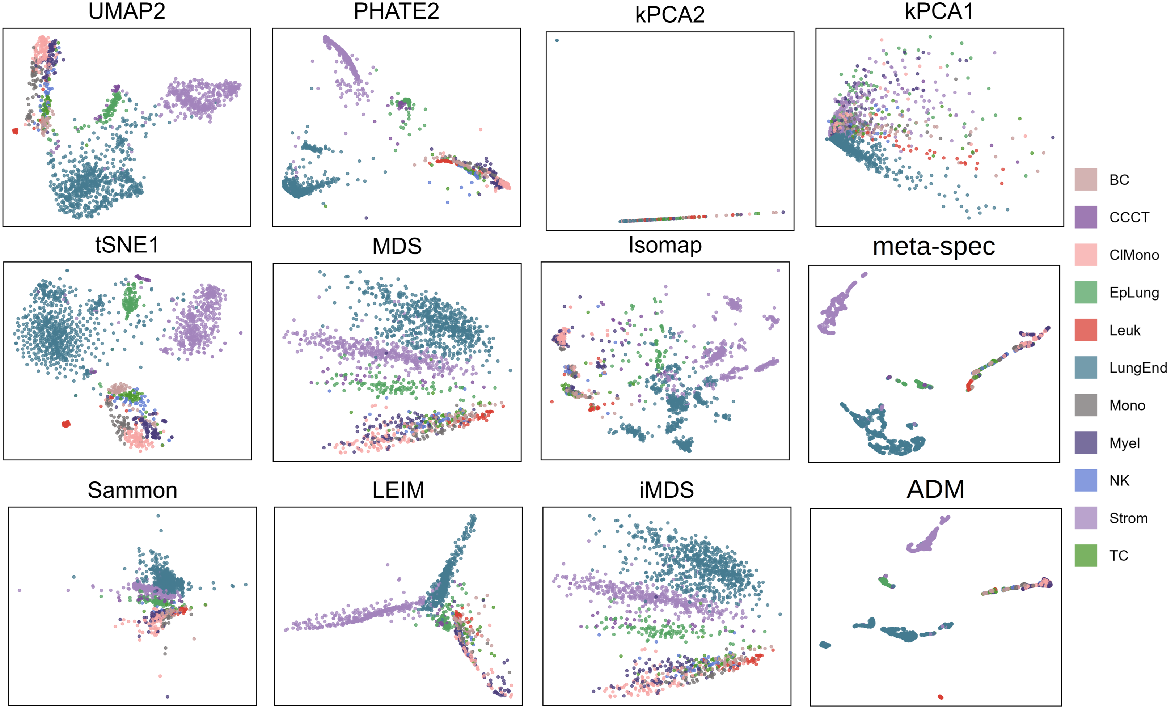
Visulization of the Quake dataset. The abbreviations in the figure correspond to their full names: BC: B cell, CCCT: ciliated columnar cell of tracheobronchial tree, ClMono: classical monocyte, EpLung: epithelial cell of lung, Leuk: leukocyte, LungEnd: lung endothelial cell, Mono: monocyte, Myel: myeloid cell, NK: natural killer cell, Strom: stromal cell, TC: T cell. The complete comparative results can be found in Figure S4 in the supplementary materials.

Compared to *Meta-Spec, ADM* exhibits more apparent separation between cell types such as B cells, lung epithelial cells, and stromal cells while maintaining higher inter-class separation and intra-class cohesion. Particularly, the distinction between B cells and T cells is more pronounced in *ADM* ‘s visualization.

Quantitative metrics, including Adjusted Rand Index (*ARI*) and Normalized Mutual Information (*NMI*), further support the performance differences between *ADM* and *Meta-Spec. ADM* demonstrates an *ARI* of 0.4224 and an *NMI* of 0.5553, whereas *Meta-Spec* exhibits an *ARI* of 0.3615 and an *NMI* of 0.5584. Additionally, *ADM* also achieves higher *CCI* scores than *Meta-Spec* on this dataset.

#### 3.3.5 CCLE dataset: cancer cell lines bulking sequencing data

The CCLE dataset [39] is a comprehensive cancer cell line resource, encompassing over 20 different types of cancer cell lines derived from a variety of tissues, including skin, central nervous system, soft tissue, ovary, blood, and lymphoid tissues. Integrating multimodal data such as gene expression, micro-RNA expression, and protein expression, CCLE offers a rich information resource for the study of cancer mechanisms, identification of biomarkers, and discovery of new therapeutic targets. In this work, we focused particularly on the gene and micro-RNA expression data provided by CCLE, analyzed as CCLE RNA and CCLE mirRNA, respectively. The CCLE RNA dataset details the expression levels of genes within different cancer cell lines, while the CCLE mirRNA dataset provides the expression profiles of micro-RNAs, using bulk RNA sequencing technique. We analyzed gene and micro-RNA expression sequencing results from 901 cell lines within CCLE, which include 14,997 genes and 700 micro-RNAs, respectively.

As shown in Figure 9, dimension-reduction and visualization based on gene and micro-RNA expression data indicated that most clustering methods effectively differentiated cancer cell lines derived from blood and lymphoid tissues from others. Meta-reduction techniques, especially *ADM*, exhibited excellent intra-group compactness, reflecting cellular similarity at the molecular level. Single dimension-reduction techniques, although capable of identifying blood and lymphoid cells, resulted in more blurred cluster boundaries and lower overall density separation.

**Fig. 9.**
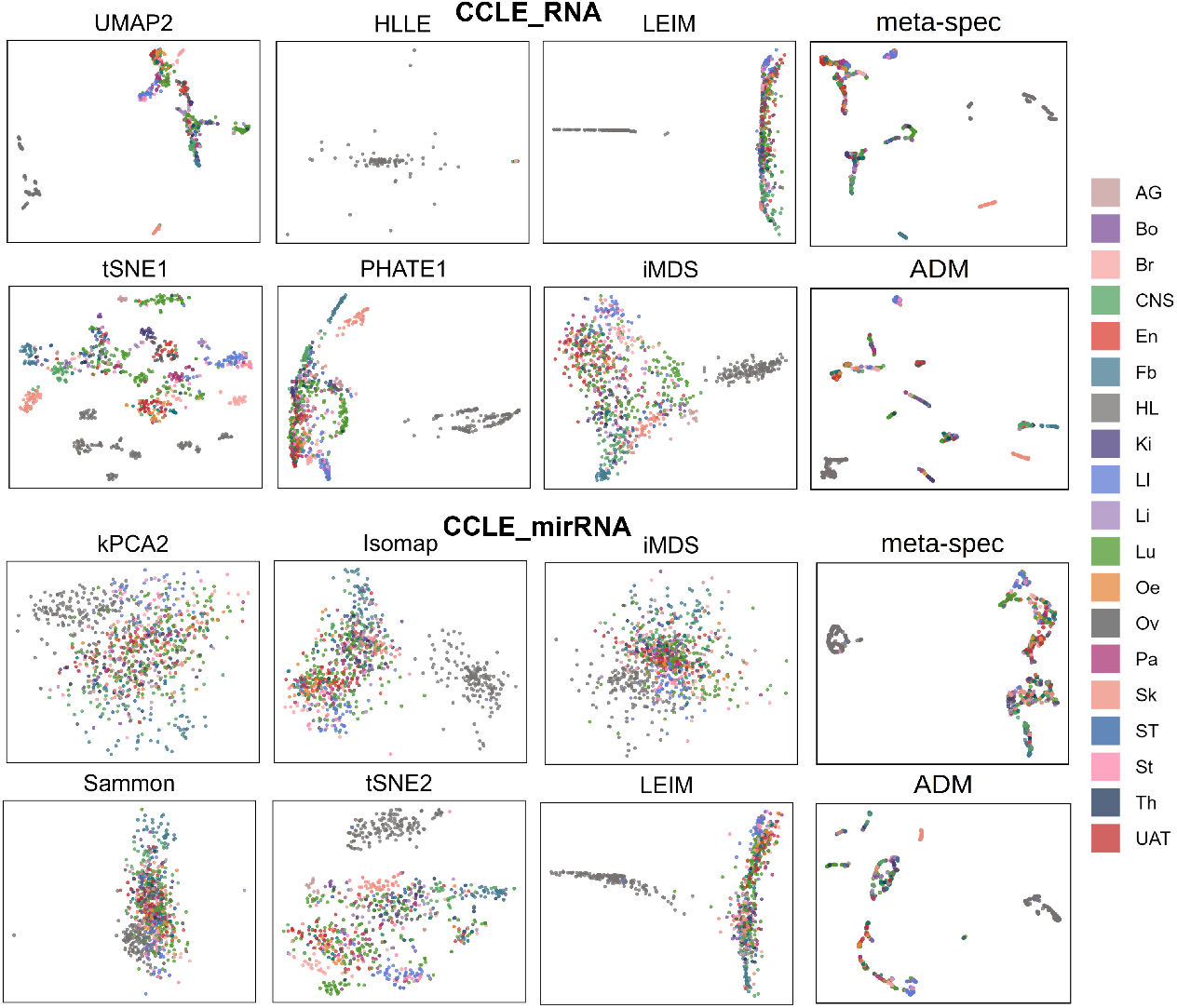
Visulization of the CCLE dataset. The abbreviations in the figure correspond to their full names: AG: Autonomic Ganglia, Bo: Bone, Br: Breast, CNS: Central Nervous System, En: Endometrium, Fb: Fibroblast, HL: Haematopoietic and Lymphoid tissue, Ki: KIDNEY, LI: Large intestine, Li: Liver, Lu: Lung, Oe: Oesophagus, Ov: Ovary, Pa: Pancreas, Sk: Skin, ST: Soft tissue, St: Stomach, Th: Thyroid, UAT: Upper Aerodigestive Tract. The complete comparative results can be found in Figures S5 and S6 in the supplementary materials.

In the visualization of central nervous system cells, the output of *ADM* is clustered into a tight group, whereas the *Meta-Spec* technique displayed more dispersed data points. *ADM* also demonstrated superior performance in clustering colon cells, whereas the *Meta-Spec* technique failed to differentiate this cell category distinctly. These observations suggest that *ADM* is more effective in maintaining intrinsic cellular similarity. Quantitatively, the *ADM* technique outperformed *Meta-Spec* for CCLE RNA, with an Adjusted Rand Index (*ARI*) of 0.3196 and a Normalized Mutual Information (*NMI*) of 0.545, compared to the *ARI* of 0.2641 and *NMI* of 0.5088 for *Meta-Spec*. for CCLE mirRNA, *ADM* ‘s *ARI* was 0.2872, and *NMI* was 0.4134, surpassing *Meta-Spec*’s *ARI* of 0.2091 and *NMI* of 0.3803.

### 3.4 Efficacy of ADM in Single-Cell Annotation

To assess the practical utility of the ADM method, we analyzed several single-cell transcriptomics datasets, which comprised various cell types with initially concealed labels. The dimension reduction and visualization were executed as outlined in Section 3.3. Subsequently, we applied the DBSCAN algorithm [40] for density-based spatial 19 clustering to delineate distinct clusters within the two-dimensional projection. We hypothesized these clusters correspond to either homogeneous cell types or similar cellular states. Differential expression analysis across these clusters was conducted using Seurat [41], providing the foundation for each cluster’s annotation.

Figure 10A and Figure 10B show that ADM highlighted the upregulated expression patterns of the MAG and APOD genes in Cluster 1, both identified as markers for oligodendrocyte cells [42–47]. This suggests that this cluster predominantly consists of oligodendrocyte cells, aligning with experimental findings from the dataset [35]. However, alternative dimension reduction methods depicted more dispersed expressions of MAG and APOD across clusters. Additionally, in Clusters 5 and 6, differential expression patterns of PLPP3 (Figure 10C) and PLTP (Figure 10D) were observed, which are markers for astrocytes [44] and endothelial cells [45], respectively. These patterns are consistent with the cell annotation labels [35].

**Fig. 10.**
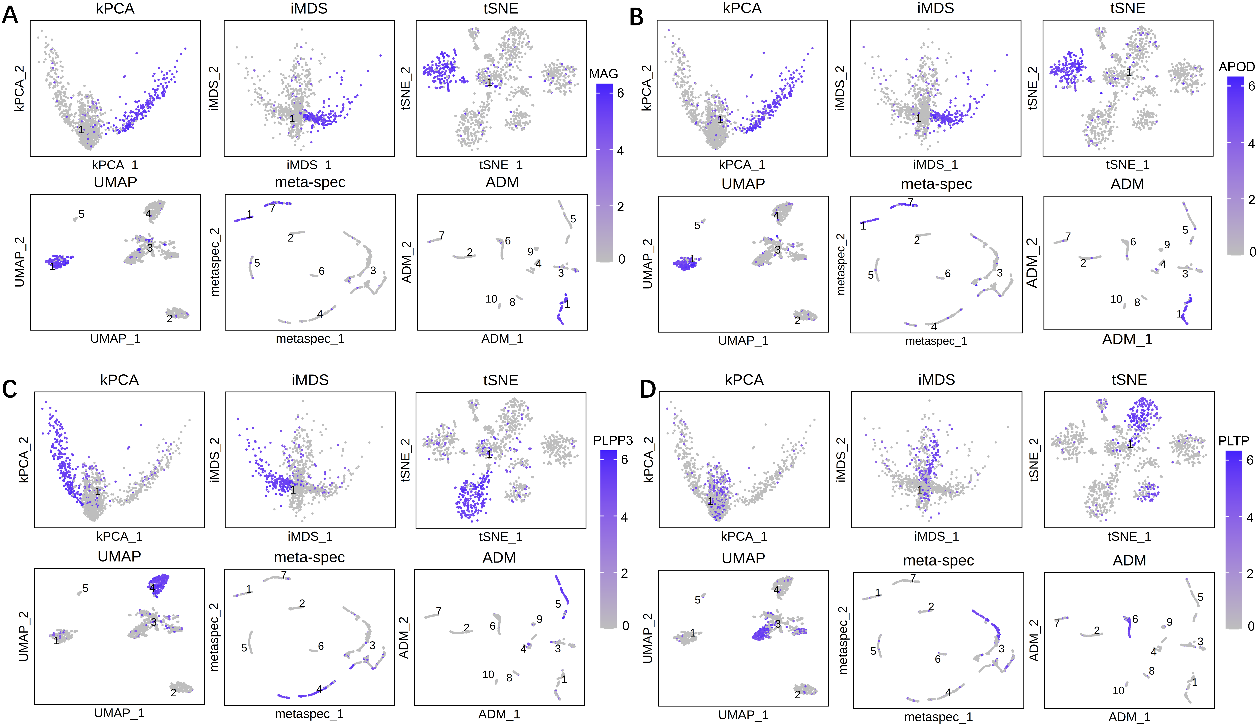
Visualization of marker genes on the dimensionality reduction results of kPCA, iMDS, t-SNE, UMAP, meta-spec, and ADM. **A:** Visualization of the distribution of MAG in the Oihane dataset. **B:** Visualization of the distribution of APOD. **C:** Visualization of the distribution of PLPP3. **D:** Visualization of the distribution of PLTP.

ADM’s ability is further corroborated by the precise alignment of key gene expression with established cellular markers, as demonstrated in the visualized clusters. This effectively validates ADM’s robust capability to perform stable dimension reduction and highlights its effectiveness in preserving the intricate structures of biological data.

## 4 Conclusion

This study introduces *ADM*, a novel meta-dimension reduction method based on dynamic Markov processes and information diffusion theory, designed to explore and integrate multiple dimension reduction results. By dynamically transforming traditional spatial metrics into diffusion distance metrics, *ADM* adapts the time scale of information diffusion based on sample-specific characteristics, enhancing the exploration of the global structure while revealing sample heterogeneity.

Extensive evaluations, including simulation studies and real-world datasets, demonstrate *ADM* ‘s superior ability to maintain data structure integrity and effectively mitigate interference from low-quality outputs. *ADM* outperforms existing single- and meta-dimension reduction methods in various assessments, offering improved visualization and a more accurate representation of the underlying data structure.

In simulations, *ADM* generally achieves higher scores in Category Consistency Index (*CCI*), Structural Consistency Index (*SCI*), Adjusted Rand Index (*ARI*), and Normalized Mutual Information (*NMI*), indicating its effectiveness in capturing and combining the strengths of various dimension reduction methods more comprehensively.

Applications on real-world single-cell datasets further validated *ADM* ‘s efficacy. It demonstrated superior performance in visualizing complex biological datasets, including those from mouse brain regions, human lymphocyte populations, mouse tendon cells, human immune and respiratory systems, and cancer cell lines. *ADM* provided clearer separation between cell types and subtypes, highlighting its capacity to enhance biological interpretations and insights.

In our future work, we intend to explore the broad potential of *ADM*, originally developed for dimension reduction and visualization, across diverse domains requiring multi-modal data integration. By facilitating the integration of disparate data sources, *ADM* is promising to offer a more comprehensive view of the data, thereby enriching the depth and breadth of analysis.

## Availability of data and materials

All code and preprocessed data used for this study are publicly available at https://github.com/Seven595/ADM. The datasets presented in this study can be acquired from the following websites or accession numbers: (1) The mouse midbrain and striatum scRNAseq data (GSE148393); (2) The human lymphocyte population dataset (GSE81772); (3) The tendon cell population dataset (GSE138515); (4) The mixed cell types of human immune and respiratory systems (GSE151334); (5) The cancer cell lines bulking sequencing data (https://depmap.org/portal/datapage/?tab=allData).

