## supplementary materials for "ADM: Adaptive Graph Diffusion for Meta-Dimension Reduction"

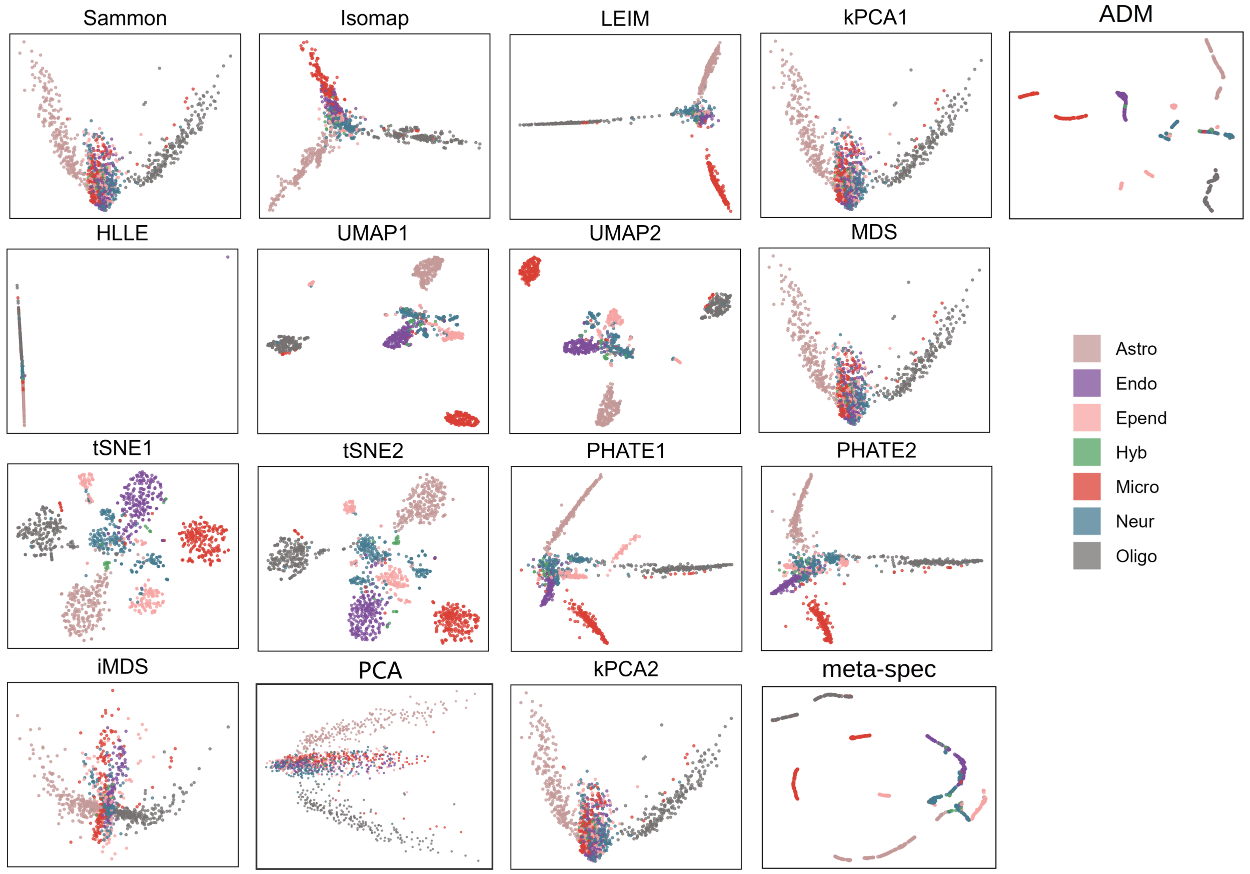


Figure S1: Visulization of the Oihane dataset on all comparative techniques. The abbreviations in the figure correspond to their full names: Astro: Astrocytes, Endo: Endothelial, Epend: Ependymal, Hyb: Hybrid, Micro: Microglia, Neur: Neurons, Oligo: Oligodendrocytes

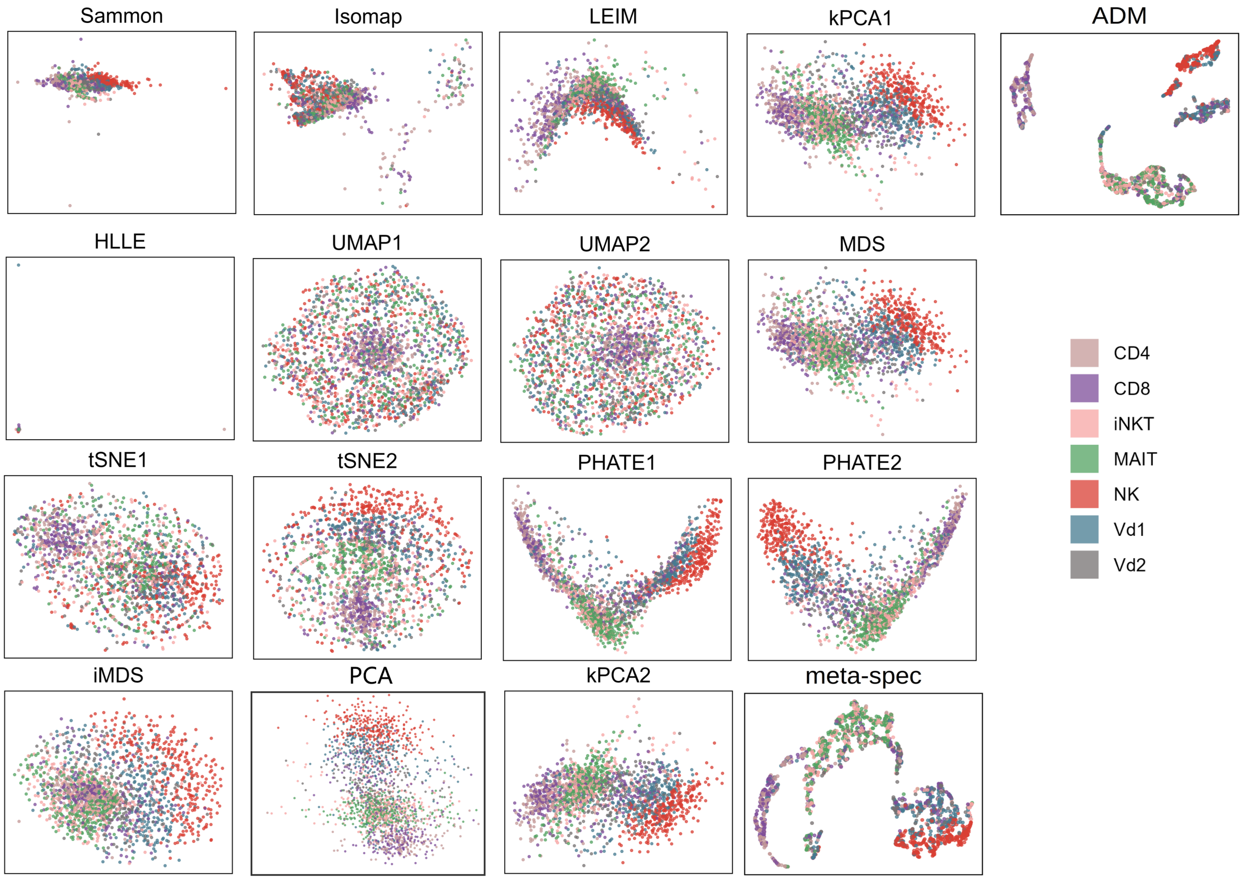


Figure S2: Visulization of Gutierrez dataset on all comparative techniques.

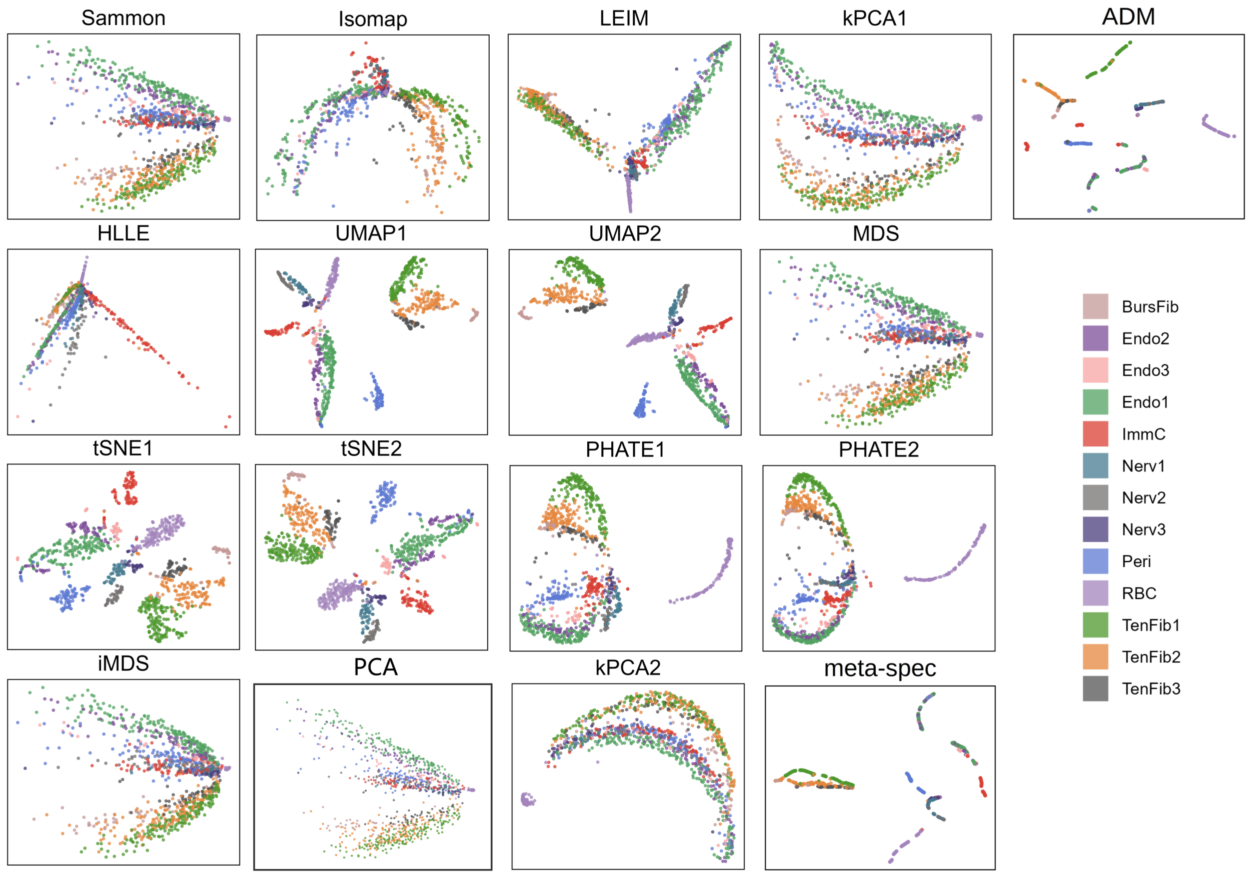


Figure S3: Visulization of the Micheli dataset. The abbreviations in the figure correspond to their full names: BursFib: Bursal fibroblasts, Endo: Endothelial cells, ImmC: Immune Cells, Nerv: Nerve cells, Peri: Pericytes, RBC: Red blood cells, TenFib: Tendon fibroblasts

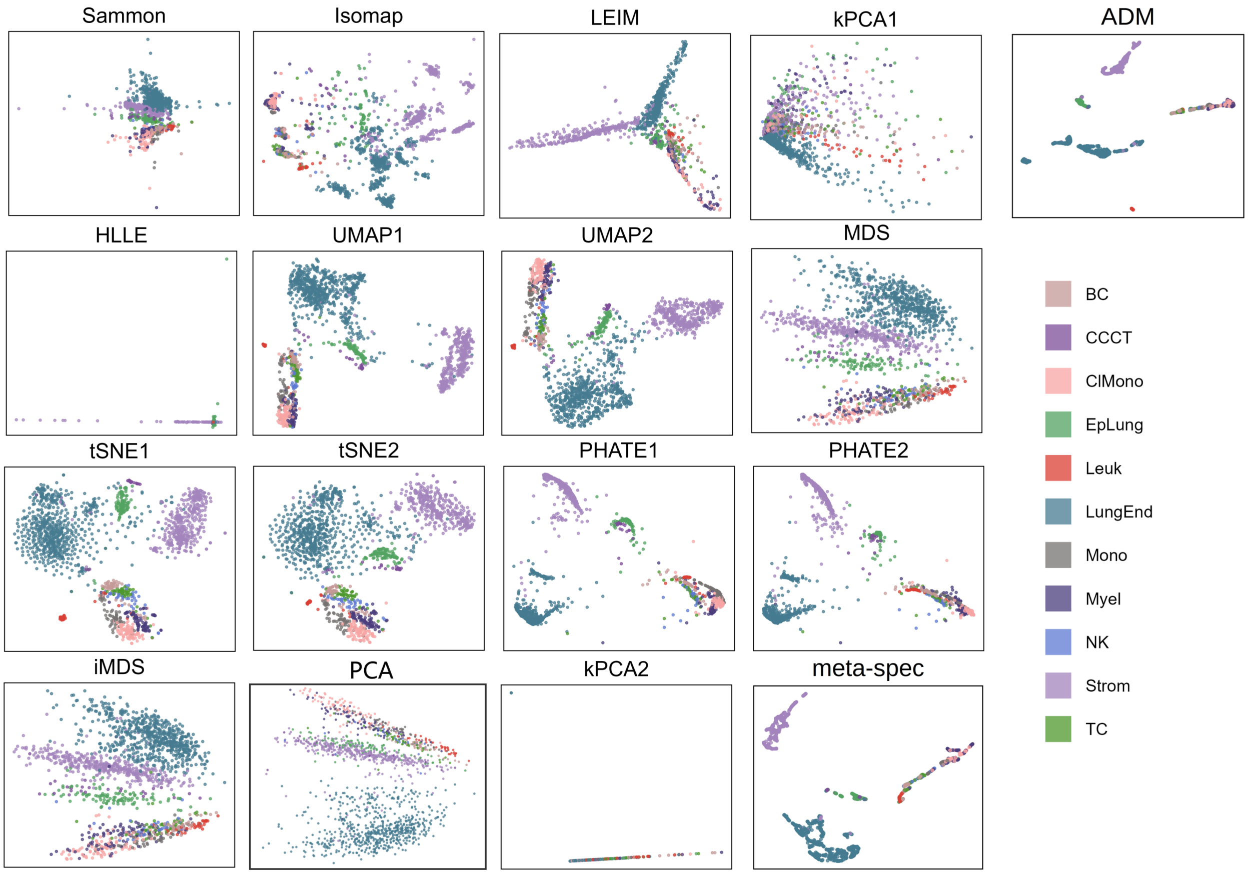


Figure S4: Visulization of the Quake dataset on all comparative techniques. The abbreviations in the figure correspond to their full names: BC: B cell, CCCT: ciliated columnar cell of tracheobronchial tree, ClMono: classical monocyte, EpLung: epithelial cell of lung, Leuk: leukocyte, LungEnd: lung endothelial cell, Mono: monocyte, Myel: myeloid cell, NK: natural killer cell, Strom: stromal cell, TC: T cell.

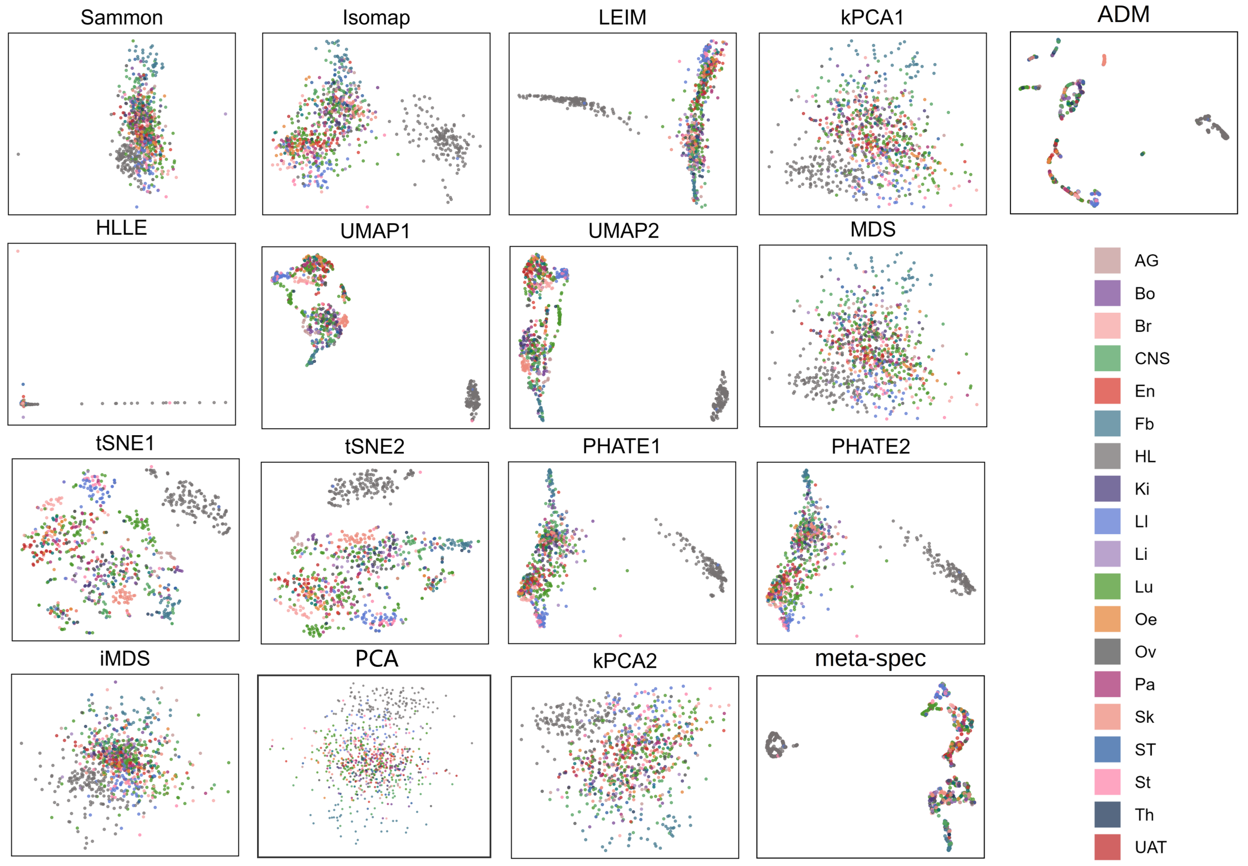


Figure S5: Visulization of the CCLE_mirRNA dataset on all comparative techniques. The abbreviations in the figure correspond to their full names: AG: Autonomic Ganglia, Bo: Bone, Br: Breast, CNS: Central Nervous System, En: Endometrium, Fb: Fibroblast, HL: Haematopoietic and Lymphoid tissue, Ki: KIDNEY, LI: Large intestine, Li: Liver, Lu: Lung, Oe: Oesophagus, Ov: Ovary, Pa: Pancreas, Sk: Skin, ST: Soft tissue, St: Stomach, Th: Thyroid, UAT: Upper Aerodigestive Tract

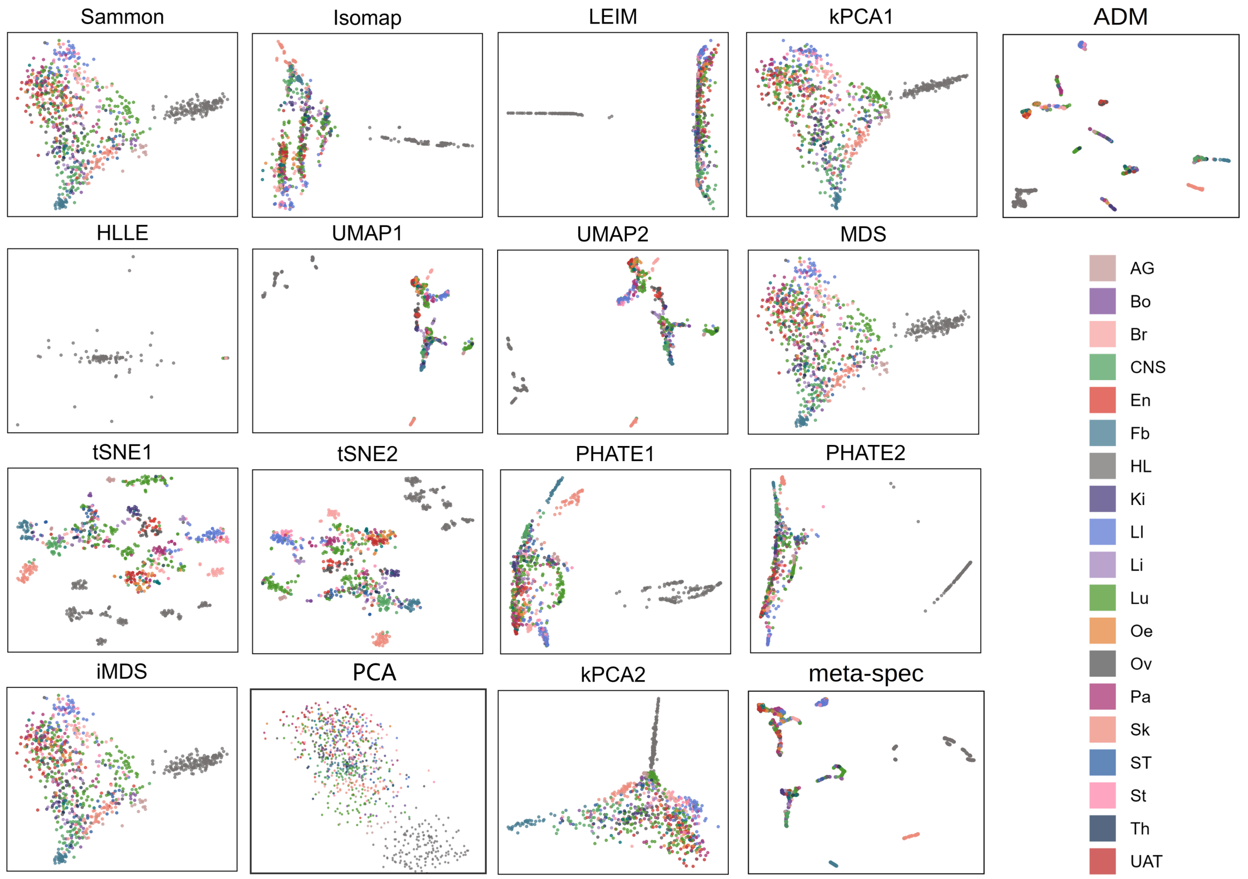


Figure S6: Visulization of the CCLE_RNA dataset on all comparative techniques. The abbreviations in the figure correspond to their full names: AG: Autonomic Ganglia, Bo: Bone, Br: Breast, CNS: Central Nervous System, En: Endometrium, Fb: Fibroblast, HL: Haematopoietic and Lymphoid tissue, Ki: KIDNEY, LI: Large intestine, Li: Liver, Lu: Lung, Oe: Oesophagus, Ov: Ovary, Pa: Pancreas, Sk: Skin, ST: Soft tissue, St: Stomach, Th: Thyroid, UAT: Upper Aerodigestive Tract
